# Spectral inference reveals principal cone-integration rules of the zebrafish inner retina

**DOI:** 10.1101/2021.08.10.455697

**Authors:** Philipp Bartel, Takeshi Yoshimatsu, Filip K Janiak, Tom Baden

## Abstract

In the vertebrate retina, bipolar cells integrate the signals from different cone types at two main sites: directly, via dendritic inputs in the outer retina, and indirectly, via axonal inputs in the inner retina. Of these, the functional wiring of the indirect route, involving diverse amacrine cell circuits, remains largely uncharted. However, because cone-photoreceptor types differ in their spectral sensitivities, insights into the total functional cone-integration logic of bipolar cell might be gained by linking spectral responses across these two populations of neurons. To explore the feasibility of such a “spectral-circuit-mapping” approach, we here recorded *in vivo* responses of bipolar cell presynaptic terminals in larval zebrafish to widefield but spectrally resolved flashes of light. We then mapped the results onto the previously established spectral sensitivity functions of the four cones.

We find that this approach could explain ∼95% of the spectral and temporal variance of bipolar cell responses by way of a simple linear model that combined weighted inputs from the cones with four stereotyped temporal components. This in turn revealed several notable integration rules of the inner retina. Overall, bipolar cells were dominated by red-cone inputs, often alongside equal sign inputs from blue- and green-cones. In contrast, UV-cone inputs were uncorrelated with those of the remaining cones. This led to a new axis of spectral opponency which was mainly set-up by red-/green-/blue-cone “Off” circuits connecting to “natively-On” UV-cone circuits in the outermost fraction of the inner plexiform layer – much as how key colour opponent circuits are established in mammals. Beyond this, and despite substantial temporal diversity that was not present in the cones, bipolar cell spectral tunings were surprisingly simple. They either approximately resembled both opponent and non-opponent spectral motifs already present in the cones or exhibited a stereotyped non-opponent broadband response. In this way, bipolar cells not only preserved the efficient spectral representations in the cones, but also diversified them to set up a total of six dominant spectral motifs which included three axes of spectral opponency. More generally, our results contribute to an emerging understanding of how retinal circuits for colour vision in ancestral cone-tetrachromats such as zebrafish may be linked to those found in mammals.

## INTRODUCTION

For colour vision, retinal circuits combine and contrast the signals from spectrally distinct types of photoreceptors^1^. For this, our own trichromatic vision uses spectral signals along two main opponent axes: “blue-yellow” and “green-red”^2–5^. Of these, blue-yellow comparisons are based on ancestral cone-type selective retinal circuits that differentially contact SWS1-(“blue”) and LWS-cones (“green/red”, aka. “yellow”), while reliably contrasting “green-red” is thought to require the central brain^1,5,6^. This is because primate “green”- and “red-cones” emerged from a relatively recent LWS gene duplication that enabled new green sensitivity in some LWS-cones, however without providing a known means for postsynaptic retinal circuits to distinguish between “green” and “red” LWS-cone variants^3,7^. Accordingly, in our own eyes, one axis of spectral opponency arises in the retina, and a second is probably decoded only in the brain.

In contrast, most non-mammalian vertebrate lineages, including fish, amphibians, reptiles, and birds, retain the full complement of ancestral cone-types based on four opsin-gene families: SWS1 (UV-cones), SWS2 (blue-cones), RH2 (green-cones), LWS (red-cones)^1,8–10^. These feed into cone-type selective circuits in the outer retina (e.g. zebrafish^11–13^, chicken^14–16^). Accordingly, in these non-mammalian lineages, the expectation is that up to tetrachromatic colour vision should be possible based on cone-opponent ancestral circuits that are developmentally hardwired into the retinal fabric, without a necessity for building additional spectral opponencies in the brain. In agreement, physiological recordings from retinal neurons in cone-tetrachromatic species including turtles^17^ and diverse species of fish^8,18–22^ consistently revealed a rich complement of complex spectral signals, including diverse spectral opponencies.

However, what the dominant opponencies are, and how they are built at the circuit level remains incompletely understood in any cone-tetrachromat vertebrate^8^. This is in part because already horizontal cells in the outer retina functionally interconnect and potentially retune cone-types^9,13,23–25^, thus limiting the possibility of making inferences about spectral processing based on recordings from downstream neurons. To address this, we recently measured the *in-vivo* spectral tuning of the synaptic outputs from the four cone-types in larval zebrafish using spatially widefield but spectrally narrow flashes of light^26^. This revealed that red-cones are non-opponent, green- and blue-cones are strongly opponent with distinct zero crossings (∼523 and ∼483 nm, respectively), and UV-cones are weakly opponent with a zero crossing at ∼450 nm. Accordingly, in larval zebrafish already the cone-output provides up to three axes of spectral opponency^8,26^. However, the opponent axis provided by UV-cones was weak, which left its role in zebrafish colour vision unclear. Moreover, in view of expected extensive mixing of cone-signals in downstream circuits^11,27^, if and how the cones’ spectral axes are propagated downstream remains unknown.

Accordingly, we asked how downstream retinal circuits make use of the spectrally complex cone signals to either consolidate or to retune their spectral axes for transmission to the brain. For this, we used two-photon (2P) imaging to measure spatially widefield but spectrally highly resolved tuning functions at the level of retinal bipolar cell (BCs) presynaptic terminals in the inner retina. This strategy was previously used to establish the spectral tunings of the cones^26,28^, thus facilitating direct comparison.

We find that all three spectral axes already set-up by the cones are conserved at the level of BC presynaptic terminals, and no new axes are created. However, the “UV-red” axis was notably boosted and diversified into numerous variants of either polarity via new opponent circuits that derive from red-/green-/blue-Off-circuits connecting to UV-On-circuits. The remaining non-opponent BCs were either broadly tuned, likely built by pooling signals from all four cone types, or essentially resembled the tunings of red- and/or UV-cones in isolation. Beyond spectral tuning, bipolar cells showed a rich complement of temporal features that were absent in cones, which were notably intermixed with spectral information.

Taken together, larval zebrafish BC-circuits for colour vision therefore directly built upon the existing cone-tunings rather than set up fundamentally new opponencies, while at the same time adding substantial temporal complexity to the retinal code.

## RESULTS

### A complex interplay of spectral and temporal signals amongst BCs

To establish *in vivo* spectral tuning functions at the level of individual presynaptic terminals of bipolar cells (BCs) in the inner retina, we imaged light-evoked calcium responses from 6-7 days post fertilisation (*dpf*) RibeyeA:SyjGCaMP7b zebrafish under two-photon (2P) using established protocols^18,29,30^ (Methods). To record from 100s of individual BC terminals in parallel, we used a non-telecentric triplane imaging approach^31^ (Methods). For light-stimulation, we used the same system and protocol previously employed to determine cone-tunings^26^ (Figure 1A,B). In brief, light from 13 spectrally distinct LEDs was collected by a collimator after reflecting off a diffraction grating which served to narrow individual LED spectra reaching the eye^32^. From here, stimuli were presented to the fish as widefield but spectrally narrow flashes of light (3 s On, 3 s Off, starting from “red” and sweeping towards UV; Methods). One example recording from BC terminals is illustrated in Figure 1C-E alongside averaged cone-responses to the same stimulus (Figure 1B) taken from Ref^26^. In short, each recording plane was automatically processed to detect the boundaries of the inner plexiform layer (IPL, Figure 1D, left) and to place regions of interest (ROIs) based on pixel-wise response coherence over consecutive repeats (Figure 1D, right, Methods). From here, fluorescence traces from each ROI were extracted, detrended, z-scored, and averaged over typically 7-8 stimulus repetitions (Figure 1D,E). This revealed a great diversity in both the spectral and the temporal composition of responses amongst BCs. For example, some ROIs were entirely non-opponent but differed in their spectral tuning and in the degree to which they “overshot” the baseline between stimulus presentations (Figure 1F, compare ROIs labelled BC1 and BC2). Other ROIs such as the one labelled BC3 were spectrally opponent, here exhibiting Off-signals to mid-wavelength stimulation but On-signals to UV-stimulation. Finally, some ROIs including the one labelled BC4 exhibited different temporal responses to long- and short-wavelength stimulation.

**Figure 1.**
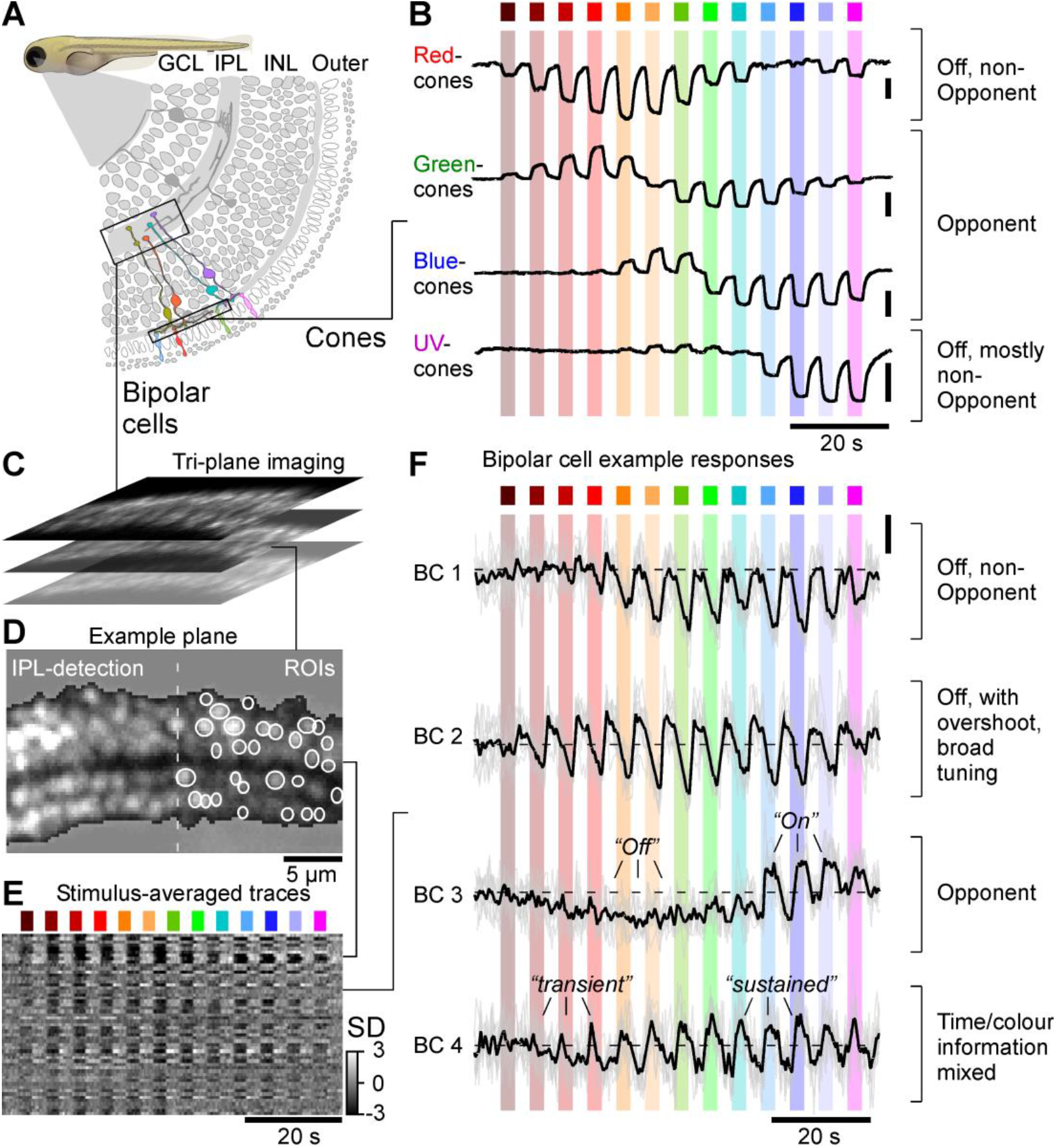
Measuring high-spectral resolution tuning curves in zebrafish bipolar cells. **A**, Schematic of the larval zebrafish retina, with cone-terminals in the outer retina and bipolar cell (BC-) terminals in the inner retina highlighted. **B**, Mean calcium-responses of red-green-, blue- and UV-cone terminals to a series of 13 spectrally distinct widefield flashes of light as indicated (data from Ref^26^). Note that for clarity the response to a 14^th^ “low-power-control” UV-LED was graphically removed compared to the original publication. **C-F**, Illustration of recording strategy for BC-terminals in the inner plexiform layer (IPL), and exemplary results. An optical tri-plane approach (C, top) was used to simultaneously record from three planes of larval zebrafish BC-terminals expressing SyGCaMP6f by way of two-photon imaging coupled with remote focussing (Methods). From here, we automatically placed regions of interest (ROIs) and detected the boundaries of the IPL (D, Methods). Time traces from all ROIs in a recording plane were z-scored and averaged across 3-5 response repeats of the full stimulus sequence (E). Example traces from individuals ROIs (F) are shown as individual repeats (grey) and averages across repeats (black).

We recorded responses from a total of n = 72 triplane scans in n = 7 fish, across four major regions of the eye: Acute Zone (AZ), Dorsal (D), Nasal (N), and Ventral (V). From here, n = 6,125 ROIs (n_AZ,D,N,V_ = 2,535, 1,172, 1,889, 529, respectively) that passed a minimum response quality criterion (Methods) were kept for further analysis.

Next, we clustered BC responses using a mixture of Gaussian model as described previously^18,21,33,34^ (Methods). This yielded 29 functional BC-clusters (Figure 2A,B), here arranged by their mean stratification position in the IPL (Figure 2C). If and how this relatively large number of functional BC-clusters maps onto veritable BC ‘types’^27^ remains unknown. For comparison, previous studies described 25 functional^18^ and 21 anatomical^11^ BCs, however a deeper census of zebrafish BC-types, for example based on additional data from connectomics^35^ and/or transcriptomics^36^ remains outstanding.

**Figure 2.**
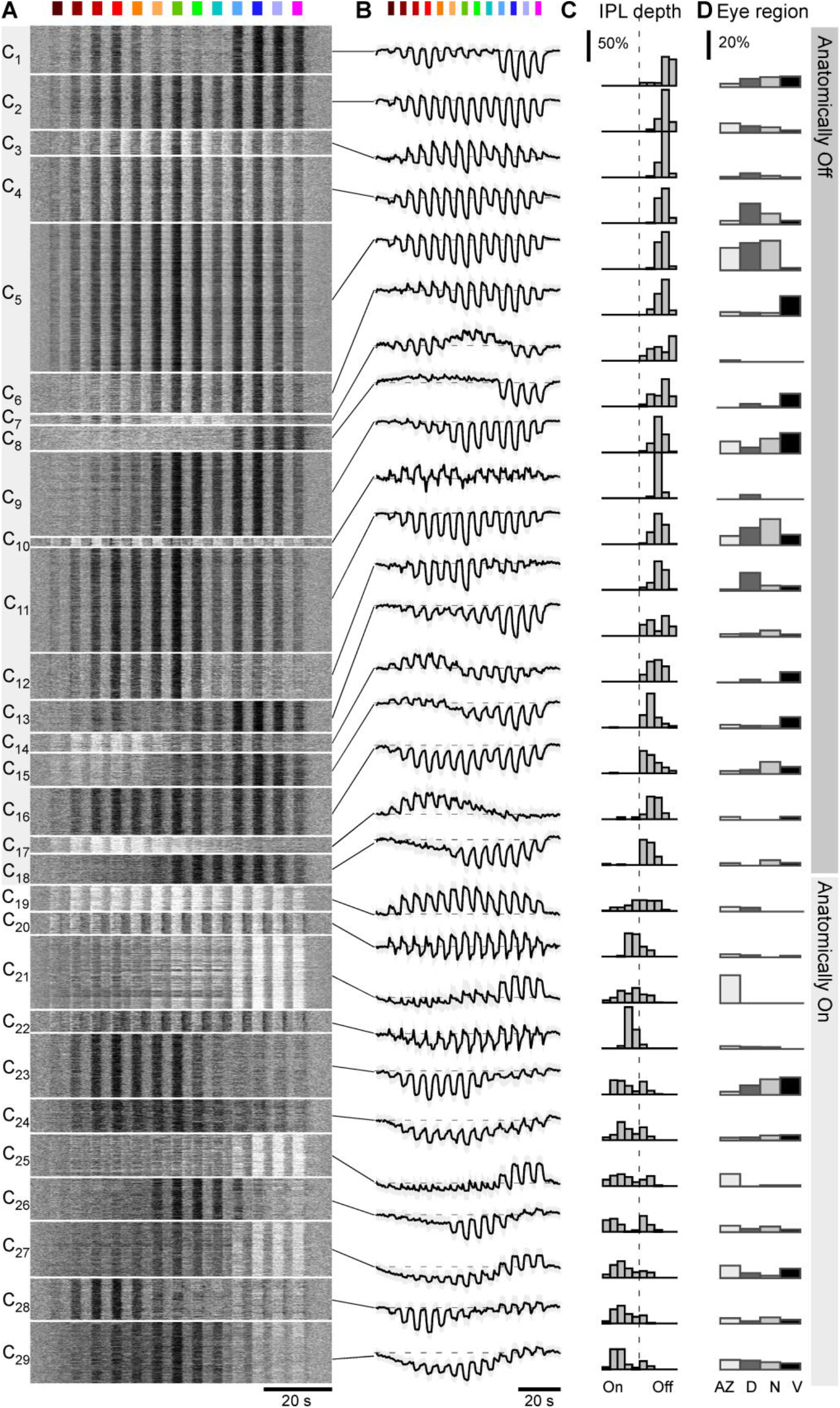
Clustering into 29 functional BC-types. **A-D**, Overview of the result from unsupervised clustering of all BC-data recorded as shown in Figure 1 that passed a minimum quality index (QI>0.4, Methods). For each cluster, shown are the individual BC-mean responses as heatmaps (A), the corresponding cluster means and SD shadings, with approximate baseline indicated in dashed (B), distribution of ROI positions in the IPL (C) and each cluster’s distribution across the four recording regions within the eye (D, from left: acute zone, dorsal, nasal, ventral). Histograms in (C) are area-normalised by cluster, and in (D) by recording region. Clusters are ordered by their average anatomical position in the IPL, starting from the border with the inner nuclear layer (cf. C).

Consistent with previous work that was based on a different stimulus with lower spectral resolution^18^, zebrafish BC-clusters were highly diverse, and many exhibited a regional bias to one or multiple parts of the eye (Figure 2D). However, with our current focus on BC-spectral tunings, we did not further analyse this eye-wide regionalisation.

Overall, BC-clusters differed strongly in their wavelength selectivity. For example, clusters C_1_ and C_2_ both hyperpolarised in response to all tested wavelengths, but C_2_ was tuned broadly while C_1_ exhibited a notable dip in response amplitudes at intermediate wavelengths. Other clusters exhibited clear spectral opponency. For example, clusters C_26-29_ all switched from Off-responses to long wavelength stimulation to On-responses at shorter wavelengths. A single cluster (C_7_) exhibited a spectrally triphasic response. BCs also differed in their temporal responses. For example, while cluster C_2_ consistently responded in a sustained manner, cluster C_3_ responses were more transient and overshot the baseline between light-flashes. Finally, diverse spectral and temporal response differences did not only exist between BC clusters, but also within. For example, cluster C_6_ switched from transient responses during long-wavelength stimulation to sustained responses during short-wavelength stimulation. In some cases, such intermixing of spectral and temporal encoding in a single functional BC-cluster could be quite complex. For example, cluster C_21_ switched from small transient On-Off responses via intermediate amplitude transient-sustained On-responses to large amplitude sustained-only On-responses in a wavelength-dependent manner.

Overall, in line with connectivity^11,37^ and previous functional work, both the spectral^18,21,22^ and the temporal diversity^18,21,22,29,38,39^ of larval zebrafish BCs long exceeded that of the cones, which at the level of presynaptic calcium were generally sustained^26^, and which only exist in four spectral variants (cf. Figure 1B).

### Linear cone-combinations using four temporal components can account for BC responses

We next explored if and how these BC cluster-means (Figure 2B) could be explained based on cone responses^26^ (Figure 3, cf. Figure 1B). For this, we implemented a simple linear model (Methods) based on the following considerations.

**Figure 3.**
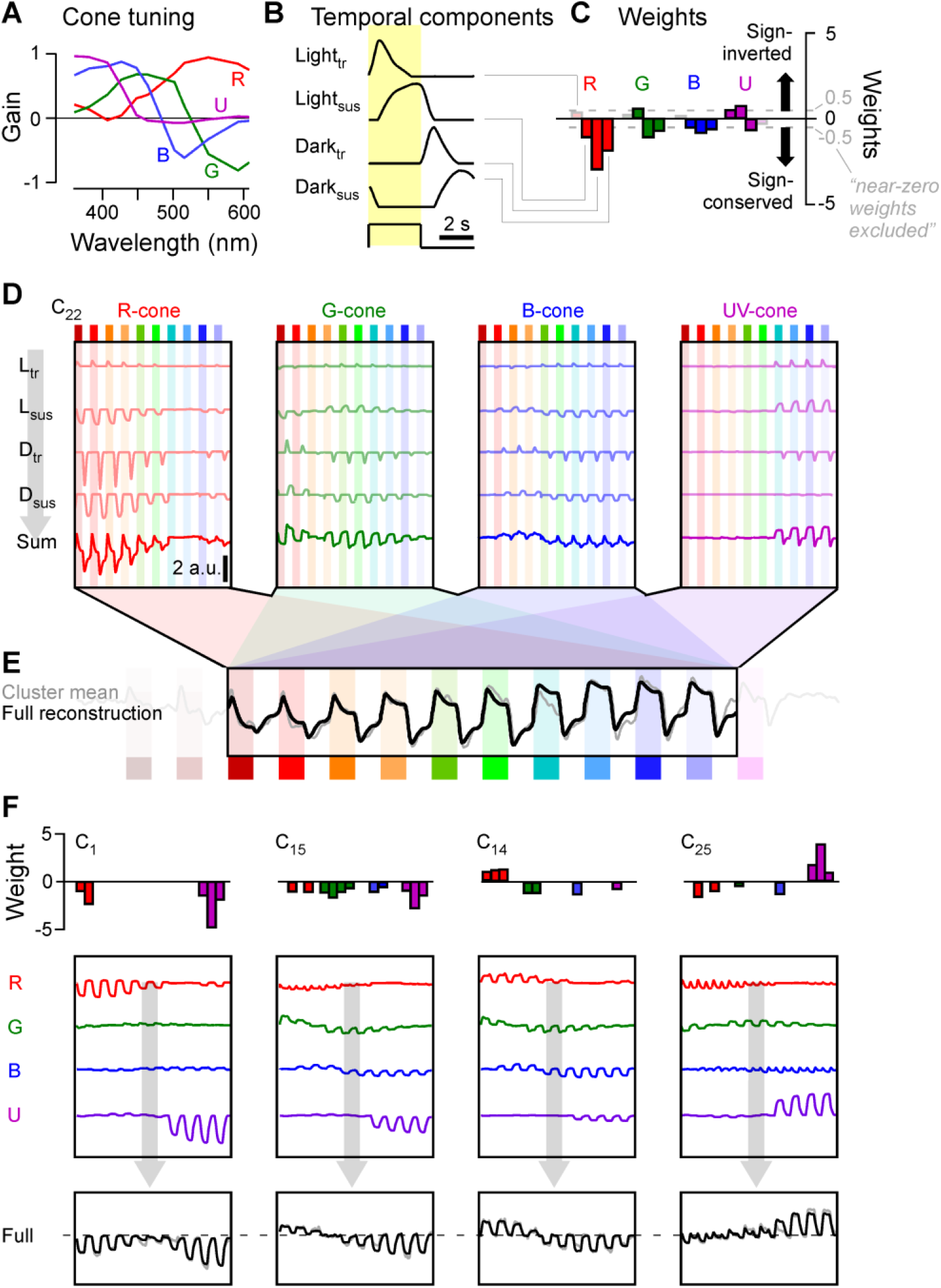
Reconstructing bipolar cell responses from cones. **A-E**, Summary of the reconstruction strategy for example cluster C_22_ (for details see Methods). Each BC-cluster reconstruction is based on the linear combination of the spectral tunings functions of the four cone-types (A, from Ref^26^) with four stereotyped temporal components associated with individual light flashes (B), yielding 4 × 4 = 16 weights (C). Weights are shown in blocks of temporal component weights (from left: Light-transient, Light-sustained, Dark-transient, Dark-sustained) associated with each cone (indicated by the corresponding colours). Bars above zero indicate sign-inverted (“On-”) weights, while bars below zero indicate sign-conserved (“Off-”) weights. The corresponding full expansion of this reconstruction is shown in (D). Individual combination of each cone’s tuning function (A) with each temporal component (B), scaled by their corresponding weight (C), yields sixteen “sub-traces” (D, upper four traces in each of the four panels, labelled L_tr_, L_sus_, D_tr_, D_sus_). Summation of each cone’s four sub-traces yields that cone’s total contribution to the cluster (D, bottom traces, labelled “sum”). Finally, summation of the four cone-totals yields the full reconstruction (E, black trace), shown superimposed on the target cluster mean (grey). **F**, as A-E, but showing only the weights (top) cone-totals (middle) and full reconstructions (bottom) for another four example clusters (from left: C_1_, C_15_, C_14_, C_25_). Further detail on reconstructions is shown in Figure S1, and all cluster’s individual results are detailed in Appendix 1.

BCs may receive cone inputs by two main, non-mutually exclusive routes: directly, via dendritic contacts onto cone-pedicles in the outer retina, and indirectly, via lateral inputs from amacrine cells in the inner retina^27^. A third route, via horizontal cells, has been proposed in the case of mice^40^. If such a route exists in zebrafish remains unknown.

In the outer retina, direct cone inputs are based on BC-type specific expression of glutamate receptor and/or transporter variants that are thought to be either all-sign-conserving or all-sign-inverting, but apparently never a mixture of both^27,38,41^. Accordingly, dendritic inputs alone should only be able to produce spectral tuning functions in BCs that can be explained by same-sign cone inputs. Any BC that cannot be explained in this manner is then expected to require spectrally distinct inputs from amacrine cells. On the other hand, variations to the temporal structure of a given cones’ contribution to a BC’s response could be implemented via either route^27,34,42^. Accordingly, we reasoned that for a linear transformation, each cone-type may feed into a functional BC-type via a unique temporal profile that represents the sum of all routes from a given cone to a given BC. In this way, our model effectively sought to explain each BC-cluster as a weighted sum of four spectral cone-tunings, but each of these four cone-inputs could have a unique temporal structure.

To capture the above considerations in a linear model, we combined the four-cone spectral tuning functions (Figure 3A, cf. Figure 1B) with four dominant temporal components extracted from BC responses: light-transient, light-sustained, dark-transient, and dark-sustained (Figure 3B, Methods). We restricted the model to capture the central ten light-stimuli (i.e. omitting the first two red-flashes and the last UV-flash) where BC-clusters generally exhibited the greatest response diversity (Figure 2).

Notably in the following paragraphs, we avoid the use of the common shorthand “On” or “Off” because in view of spectral opponency already present in cones^26^ a sign-conserving input to a BC is not categorically “Off”, and vice versa a sign-inverting input is not categorically “On”. Instead, we use the terms “light” and “dark” response, in reference to a response that occurs in the presence or absence of a light-stimulus, respectively. Also note that all extracted spectral tuning functions (e.g. Figure 3A) are x-inverted compared to the time-axes in recordings and reconstructions (e.g. Figure 3D,E). This was done because recordings were performed from long-to short wavelength stimuli, but spectral tuning functions are conventionally plotted from short-to long-wavelengths. Weights were scaled such that the mean of their magnitude equalled one, with weights <0.5 (“near-zero”) excluded from the summary plots for visual clarity. Full weights, including a detailed overview of each cluster, are available in Appendix 1.

Figures 3C-E illustrate the intermediate steps (Figure 3C,D) and final output (Figure 3E) of the model for example cluster C_22_. This functional BC-type was broadly tuned but switched from transient responses to long wavelength stimulation to more sustained responses at shorter wavelengths (Figure 3E, grey trace, cf. Figure 2A,B). To capture this behaviour (Figure 3E, black trace), the model drew on all four cones (Figure 3C), however with a particularly strong sign-conserved contribution from red-cones (Figure 3C, left). Here, the model placed a strong sign-conserving weight onto the dark-transient (D_tr_) component of the red-cone (Figure 3D, left, third trace). The strength and sign of this weight is illustrated in Figure 1C (third downwards facing red bar). In addition, the model also placed weaker sign-conserving weights onto the dark-sustained (Figure 3D, left, fourth trace) and light-sustained (second trace) components, and a weak sign-inverted weight onto the dark-transient component (first trace). Summation of these four kinetic components yielded the total modelled red-cone contribution to this cluster (Figure 3D, bottom trace).

The same principle was applied across the remaining three cones, yielding a total of sixteen (four cones times four temporal components) weights per cluster (cf. Figure 3C). In the example presented, weights were mostly sign-conserving (facing downwards). However, to capture the relatively complex temporal dynamics of this cluster, which systematically overshot the baseline between flashes, the model also drew on a number of weaker sign-inverted weights (facing upwards), for example for all light-transient components.

Figure 3F illustrates outputs of the model for another four example clusters with diverse spectral and temporal behaviours. Of these, the spectrally bimodal but “temporally simple” response profile of C_1_ was well-approximated by all sign-conserving inputs from red- and UV-cones (Figure 3F, left). Similarly, the spectrally opponent behaviour of C_15_ could be captured by all-sign-conserving inputs from all four cones (Figure 3F, second panel). Accordingly, as expected from the cone-tunings, generating opponent responses at the level of BC terminals does not categorically require new sign-opposition in the inner retina – instead, the opponency can simply be inherited from the cones. Nevertheless, not all opponent BC responses could be explained in this manner. For example, opponent cluster C_14_ required sign-inverted inputs from red-cones but sign-conserving inputs from green-, blue- and UV-cones (Figure 3F, third panel). Finally, even the more complex spectral and temporal BC-clusters could be well-approximated by relatively simple cone-mixtures. For example, C_25_ was captured by combining sign-conserved light- and dark-transient inputs from red- and blue-cones with mostly sustained and sign-inverted inputs from UV-cones (Figure 3F, rightmost).

**Figure S1 - related to Figure 3.**
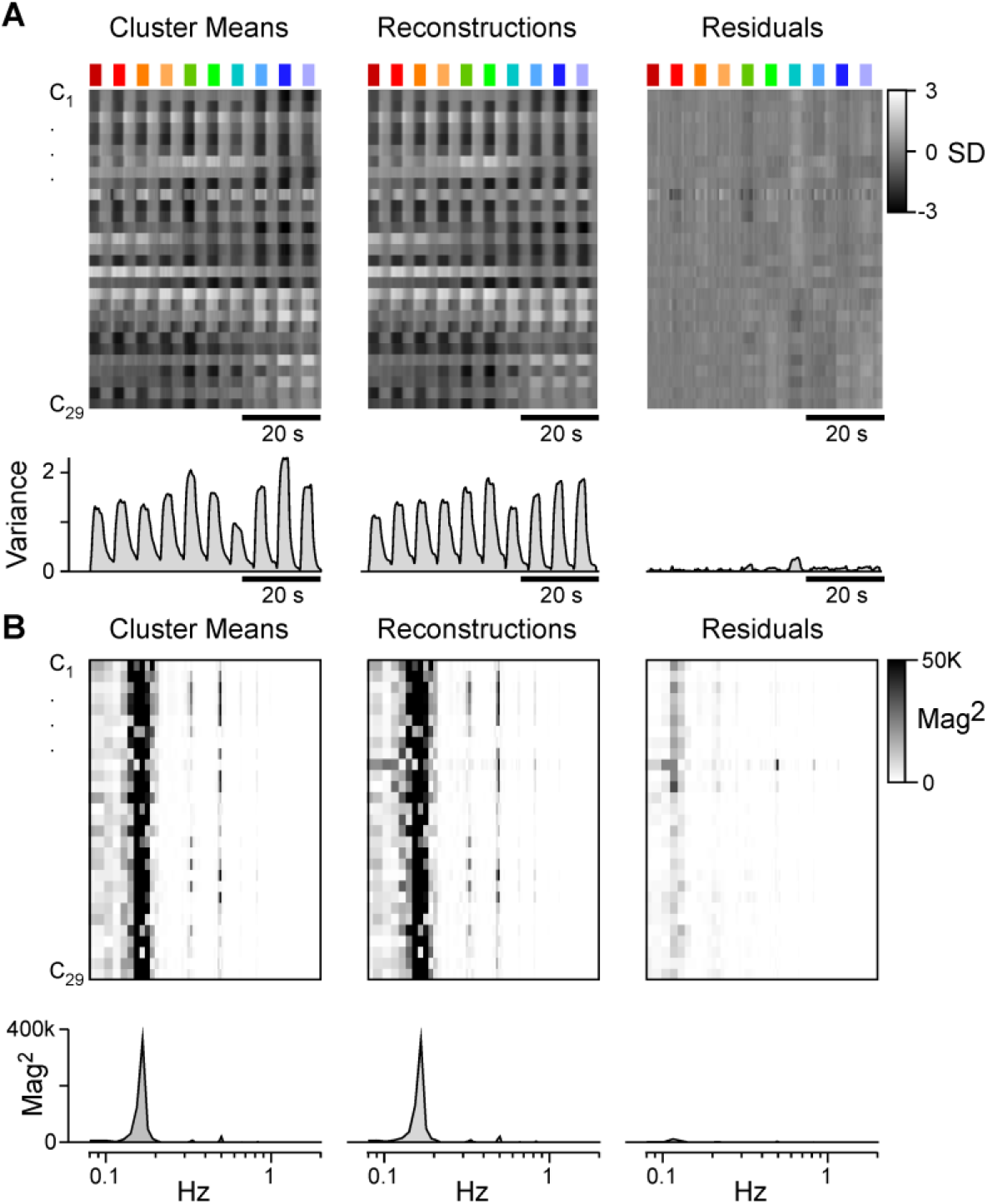
Cluster reconstruction details. **A**, Time-aligned heatmaps of all cluster means (left) are shown alongside their corresponding reconstructions (middle) and residuals (right). The time trace below each cluster shows the total variance across all clusters per time point (Methods). **B**, as A, but for magnitude-squared Fourier transforms of each cluster, reconstruction, and residuals. The traces below each panel show the averages of these transforms across all clusters (Methods). Note that for both (A) and (B), residuals retain only a small fraction of the original signal, indicating high reconstruction fidelity. Reconstruction quality of each individual cluster can further be assessed in Appendix 1.

Overall, this linear fitting procedure captured ∼95% of the total variance across the 29 cluster means (Figure S1A, Methods). Similarly, the fits also captured ∼95% of the temporal detail, based on comparison of the mean power spectra of the cluster means and that of the residuals (Figure S1B, Methods). The full result of this process is summarised in Figure 4, each time showing the cluster mean (grey) and reconstruction (black) alongside weight-summaries per cone following the schema illustrated in Figure 3B,C. Further detail is shown in Appendix 1.

**Figure 4.**
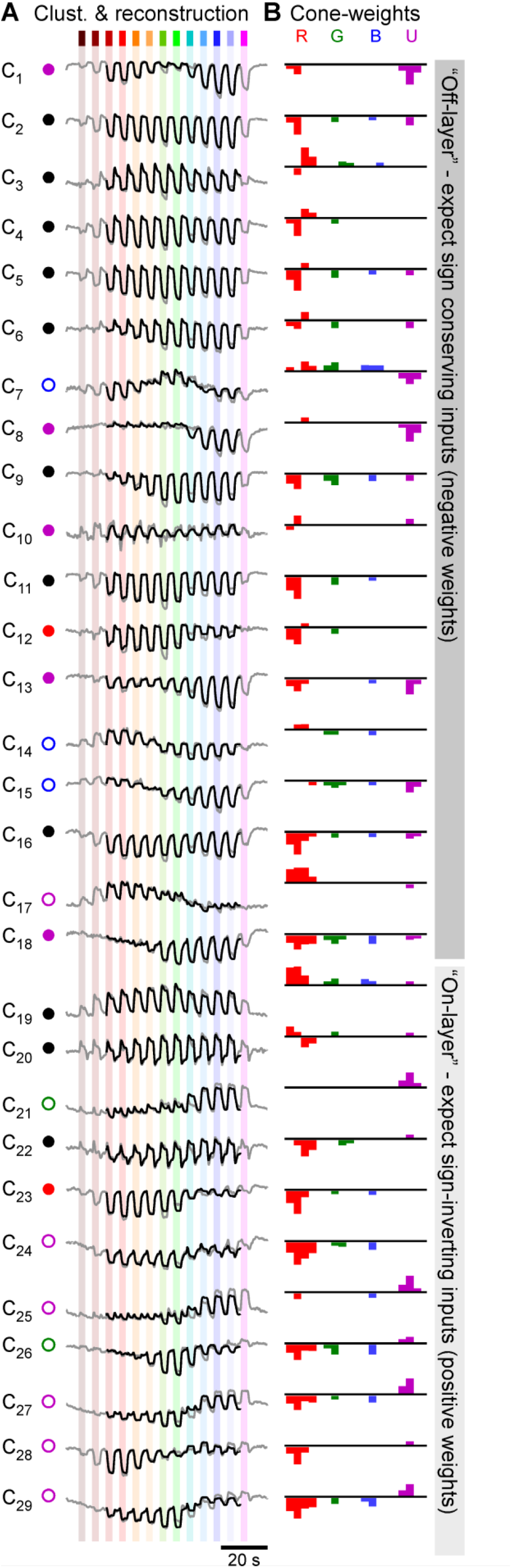
A functional overview of cone-bipolar cell mappings. **A**,**B**, Overview of all BC-cluster means (A, grey traces, cf. Figure 2B) and their full reconstructions based on the strategy detailed in Figure 3 (black traces). Associated weights are shown in (B). For clarity, “near-zero” weights (abs(w)<0.5) are omitted. Full weights are shown in Appendix 1. Note that based on outer retinal inputs only, weights are generally expected to be sign-conserving for clusters in the traditional “Off” layer (C_1_-C_18_), and sign-inverting in the anatomical “On” layer (C_19_-C_29_), as indicated on the right. The round symbols plotted next to each cluster (A) denote their allocated spectral group, as detailed in Figure 6 and associated text.

Based on the traditional separation of the inner retina into “Off-” and “On-layers”^27^, we may correspondingly expect mainly sign-conserving (negative) weights in “Off-stratifying” clusters C_1-_C_18_, and mainly sign-inverting (positive) weights for “On-stratifying” clusters C_19_-C_29_. However, this expectation was not met in several cases, for example for most of the On-stratifying clusters which nevertheless showed a general abundance of negative (“Off”) weights for red-, green- and blue-cone inputs. From here, we next explored the general rules that govern overall cone-signal integration by BCs.

### The inner retina is dominated by red-cone inputs

First, we computed histograms of all weights per cone (Figure 5A) and per temporal component (Figure 5B) to determine the dominant input-motifs across the population of all BCs. This revealed that overall, the amplitudes of red-cone weights tended to be larger than those of all other cones (red absolute weights W_R_ = 1.82±1.22; W_G,B,U_ = 0.68±0.47, 0.62±0.45, 0.87±0.88, respectively, range in SD; p<0.001 for all red-combinations, Wilcoxon Rank Sum Test). Similarly, light-response component weights tended to be larger than dark-response component weights (W_LT, LS, DT, DS_ = 0.94±0.75, 1.73±1.20, 0.85±0.8, 0.48±0.54, respectively Figure 5B). Here, the light-sustained response components that already dominate the cones (cf. Figure 1B) remained largest overall also in BCs (p<0.001 for all Light_sus_-combinations, Wilcoxon Rank Sum Test).

**Figure 5.**
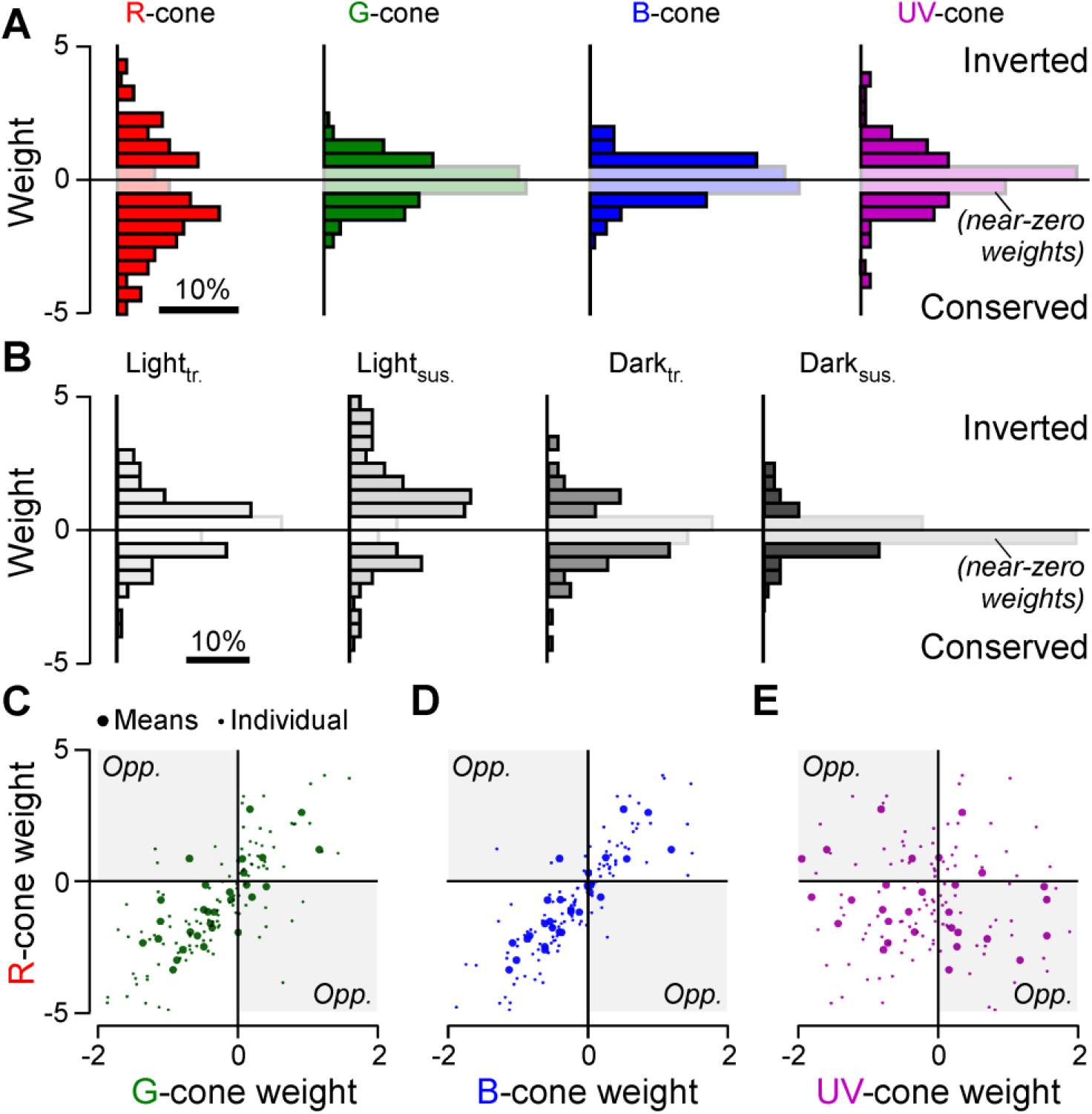
Major trends in the distribution of cone-weights. **A**,**B**, Histograms of all weights associated with inputs to each of the four cones across all clusters, independent of temporal-component types (A), and correspondingly histograms of all weights associated with temporal components, independent of cone-type (B). “Near-zero” weights (abs(w)<0.5) are graphically de-emphasised for clarity. All weights contributed equally to these histograms, independent of the size of their corresponding cluster. **C-E**, Scatterplots of all clusters’ weights associated with each cone plotted against each other as indicated. Large symbols denote the mean weight associated with each cone and cluster across all four temporal components (i.e. one symbol per cluster), while small symbols denote each weight individually (i.e. four symbols per cluster, corresponding to L_tr_, L_sus_, D_tr_, D_sus_). The remaining three possible cone-correspondences (G:B, G:U, B:U) are shown in Figure S2.

### Red-, green- and blue-cone weights co-vary independent of UV-cone weights

Next, we explored the weight relationships between the four cone types across clusters. In general, a strong correlation between weights attributed to any two cone types would suggest that inputs from these cones tend to be pooled, for example by the dendrites of individual BCs contacting both cone-types. In contrast, a low correlation or even anticorrelation between cone-weights could indicate the presence of cone-opponency.

Across clusters, we found that red-cone weights strongly correlated the weights of both green-(ρ = 0.73; 95% confidence intervals (CI) 0.49/0.86, Figure 5C) and blue-cones (ρ = 0.87, CI 0.74/0.94, Figure 5D; green vs. blue: ρ = 0.89; CI 0.77/0.95, cf. Figure S2A). The tight association between red-, green- and blue-cone weights extended across both the all-sign inverting (bottom left) and the all-sign-conserving (top right) quadrants and comprised few exceptions in the two remaining quadrants that would indicate cone-opponency. Accordingly, zebrafish BCs did not tend to differentially combine inputs from red-, green- or blue-cones of either polarity to set up potentially new opponent-axes.

In contrast, red-cone weights were uncorrelated with UV-cone weights (ρ = -0.21, CI -0.55/0.14, Figure 5E, green sc. UV: ρ = -0.04, CI -0.40/0.34; blue vs. UV; ρ = -0.34, CI -0.63/0.03, see Figure S2B,C), with many clusters scattering across the two sign-opponent quadrants (i.e. top left, bottom right). Accordingly, reconstructing a substantial fraction of BC clusters required opposite sign inputs from red-/green/blue-versus UV-cones, suggestive of a newly set-up form of spectral opponency in the inner retina. Accordingly, we next explored the spectral tuning of BC-clusters in further detail.

**Figure S2 - related to Figure 5.**
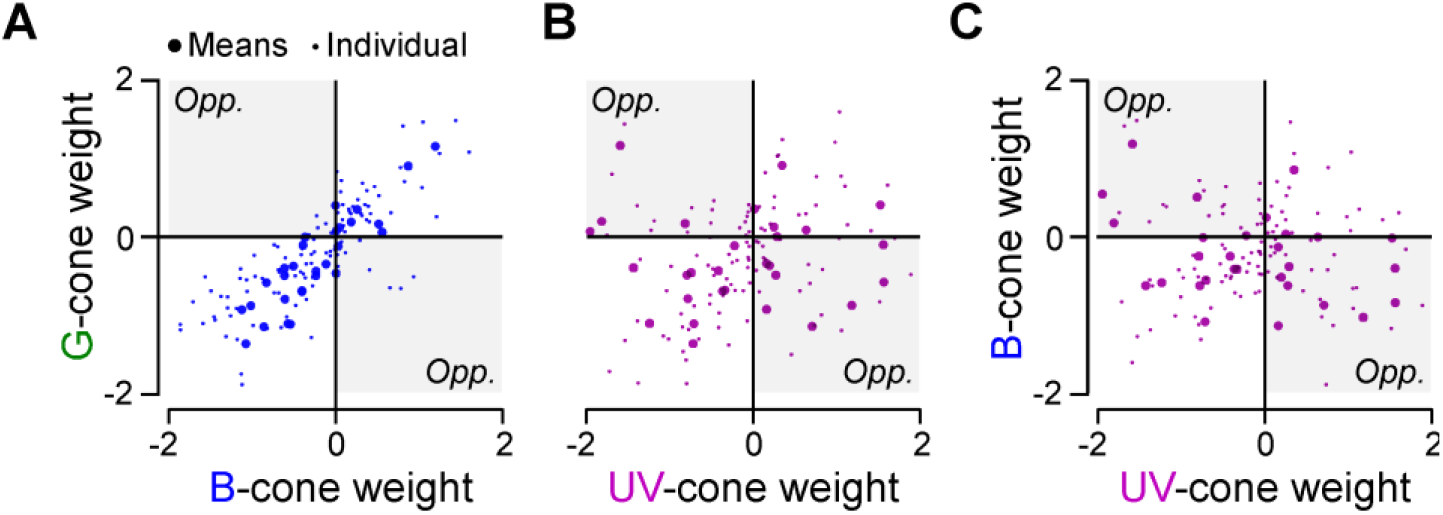
Additional cone-weight relationships. **A-C**, As Figure 5C-E, but showing weight correspondences between green-blue, green-UV and blue-UV cones, respectively.

### BC spectral responses fall into three opponent, and three non-opponent groups

The complex interplay of temporal and spectral structure in BC-responses (Figure 2) meant that their spectral tuning functions could not easily be extracted directly from the BC-cluster means, for example by means of taking the area under the curve in response to each flash of light. Instead, we estimated their tuning functions based on their fitted cone-weights (cf. Figure 4). To this end, for each cluster we summed sixteen cone-tuning functions (based on Figure 3A), each scaled by the cluster’s associated sixteen weights (i.e. red-L_tr_ + red-L_sus_.+ red-D_tr._ and so on). This summarised each cluster’s ‘bulk’ response in a single spectral tuning function that gave equal weight to each of the four temporal components (Figure 6A-F). By this measure, 18 of the 29 BC-clusters were non-opponent (62%, Figure 6A-C) and 11 were opponent (38%, Figure 6D-F). Here, opponency was defined as any tuning function that crossed and overshot zero at least once with an amplitude of at least 10% compared to that of the opposite (dominant) polarity peak response.

**Figure 6.**
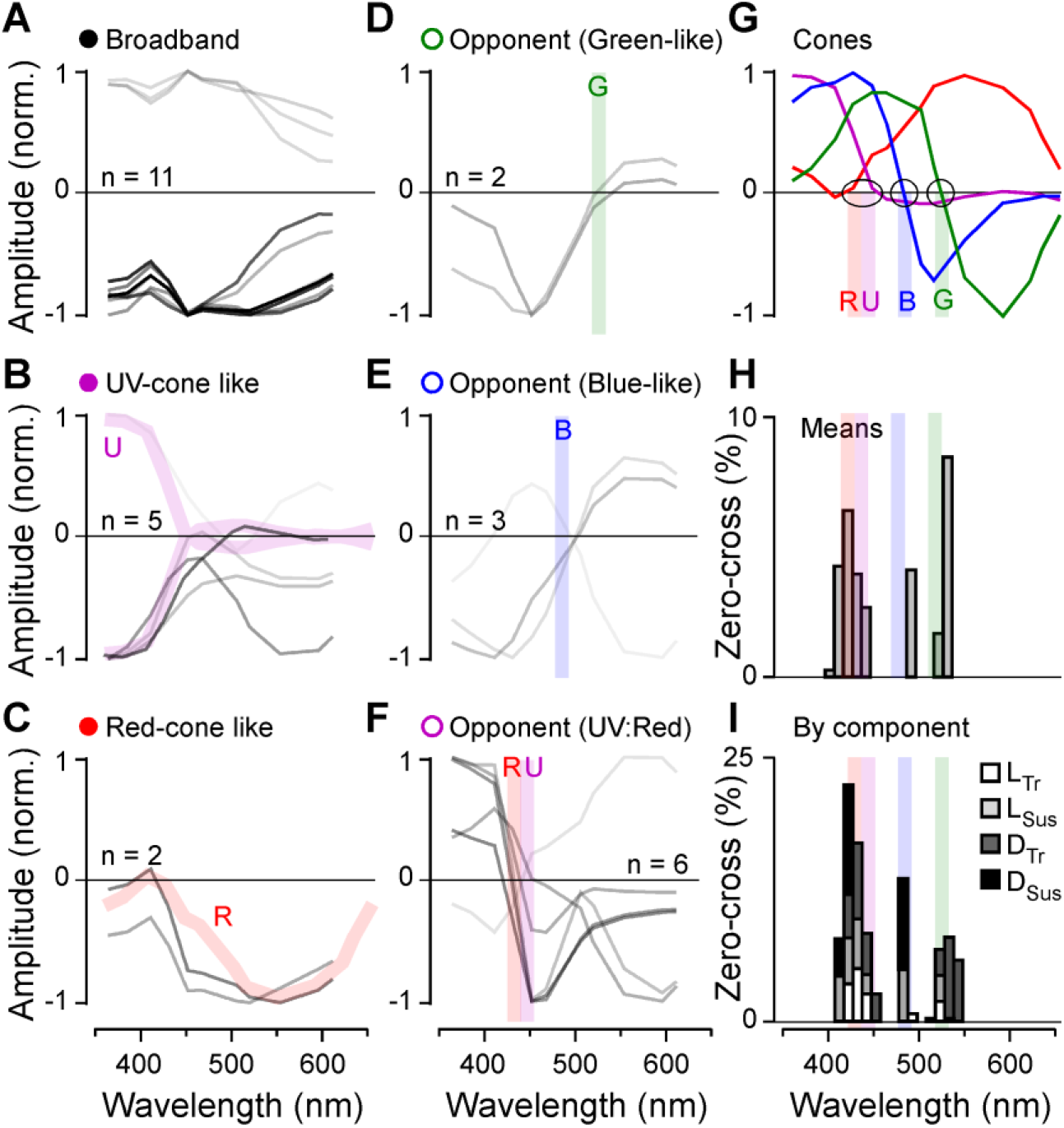
Spectral tuning of functional BC-types. **A-F**, Peak-normalised ‘bulk’ spectral tuning functions of all 29 clusters, grouped into six categories as indicated. The strength of each line indicates the numerical abundance of ROIs belonging to each cluster (darker shading = larger number of ROIs; exact number of ROIs contributing to each cluster are listed in Appendix 1). As appropriate, spectral tuning functions of cones (cf. G) are shaded into the background, as appropriate (B,C, thick coloured traces) to illustrate the close spectral correspondences of associated cones and BCs. Similarly, for three spectrally opponent groups (D-E), the approximate positions of the corresponding cone’s zero crossings are indicated with a vertical shaded line (cf. G). **G**, Cones’ spectral tuning functions, with approximate zero-crossings (blue-/green-cones) and zero-positions (red-/UV-cones) graphically indicated. **H, I**, Histograms of zero-crossings across all BC-clusters, incorporating the abundance of ROIs belonging to each cluster. Shown are crossings of ‘bulk’ spectral tunings functions (H, cf. A-F), and of spectral tuning functions that were computed each temporal component individually, as indicated (see also Figure S3C-E, and Appendix 1). Note the three prominent peaks of zero-crossing positions, approximately aligned with the zero-positions/crossings of the cones. These peaks largely disappeared when time-components were permuted at random across cones (Figure S3A,B).

Non-opponent clusters (‘closed’ symbols, cf. Figure 4A) approximately adhered to three major groups: spectrally broad (three On- and eight Off-clusters, Figure 6A), approximately UV-cone-like (one On- and four Off-clusters, Figure 6B), and approximately red-cone-like (two Off-clusters, Figure 6C). For simplicity, non-opponent clusters that combined both red- and UV inputs (e.g. C_1_) were included in the UV-group (Figure 6B), because UV-input was generally dominant over red in these cases.

Next, like non-opponent clusters, also opponent clusters (‘open’ symbols) fell into three major groups based on the spectral positions of their zero crossings: Two green-cone-like clusters (both short_Off_/long_On_, crossing at 520 and 536 nm, Figure 6D), three blue-cone-like clusters (two short_Off_/long_On_ crossing at 497 and 499 nm, plus the single triphasic C_7_ with a dominant short_On_/long_Off_ zero crossing at 490 nm, Figure 6E), and six UV-cone versus red-/green-/blue-cone opponent clusters (henceforth: UV:R/G/B, five short_On_/long_Off,:_ crossing at 416, 425, 428, 435, 448 nm, one short_Off_/long_On_ crossing at 438 nm, Figure 6F). In comparison, green- and blue-cone zero-crossings, respectively (Figure 6G, from Ref^26^) occurred at ∼523 and ∼483 nm, while red- and UV-cones, respectively, approached zero between ∼425 and 450 nm (Figure 6D-I, shadings).

The tight correspondence between opponent BC-clusters (Figure 6D-F) and cone-tunings (Figure 6G) was further illustrated by the histogram of BC-zero-crossings that also incorporated relative abundances of ROIs contributing to each cluster (Figure 6H). The histogram showed three clear peaks that were well-aligned to the three spectral axes set-up in the cones (shadings). Further, the histogram also retained its overall shape when the four temporal components underpinning each cluster were considered individually (Figure 6I). As a control, this trimodal structure largely disappeared when temporal-components were randomly shuffled between cones (Figure S3A,B), suggesting that the measured BC tunings emerged from non-random effective cone-inputs. In support, and despite appreciable diversity, the spectral tuning functions of the four temporal components that contributed to a given cluster tended to be positively correlated among both opponent and non-opponent clusters (Figure S3C-F).

Remarkably therefore, it appears that by and large, BCs tended to retain many of the dominant spectral properties of the cones rather than build fundamentally new spectral axes – all despite integrating across multiple cone types and presumably diverse inputs from spectrally complex ACs^22^. The only two notable deviations from this observation were a highly stereotypical spectral broadening in 11 clusters (Figure 6A), which may be linked to outer retinal cone-pooling^11^, and, strikingly, the emergence of six strongly UV:R/G/B opponent clusters (Figure 6F).

**Figure S3 - related to Figure 5.**
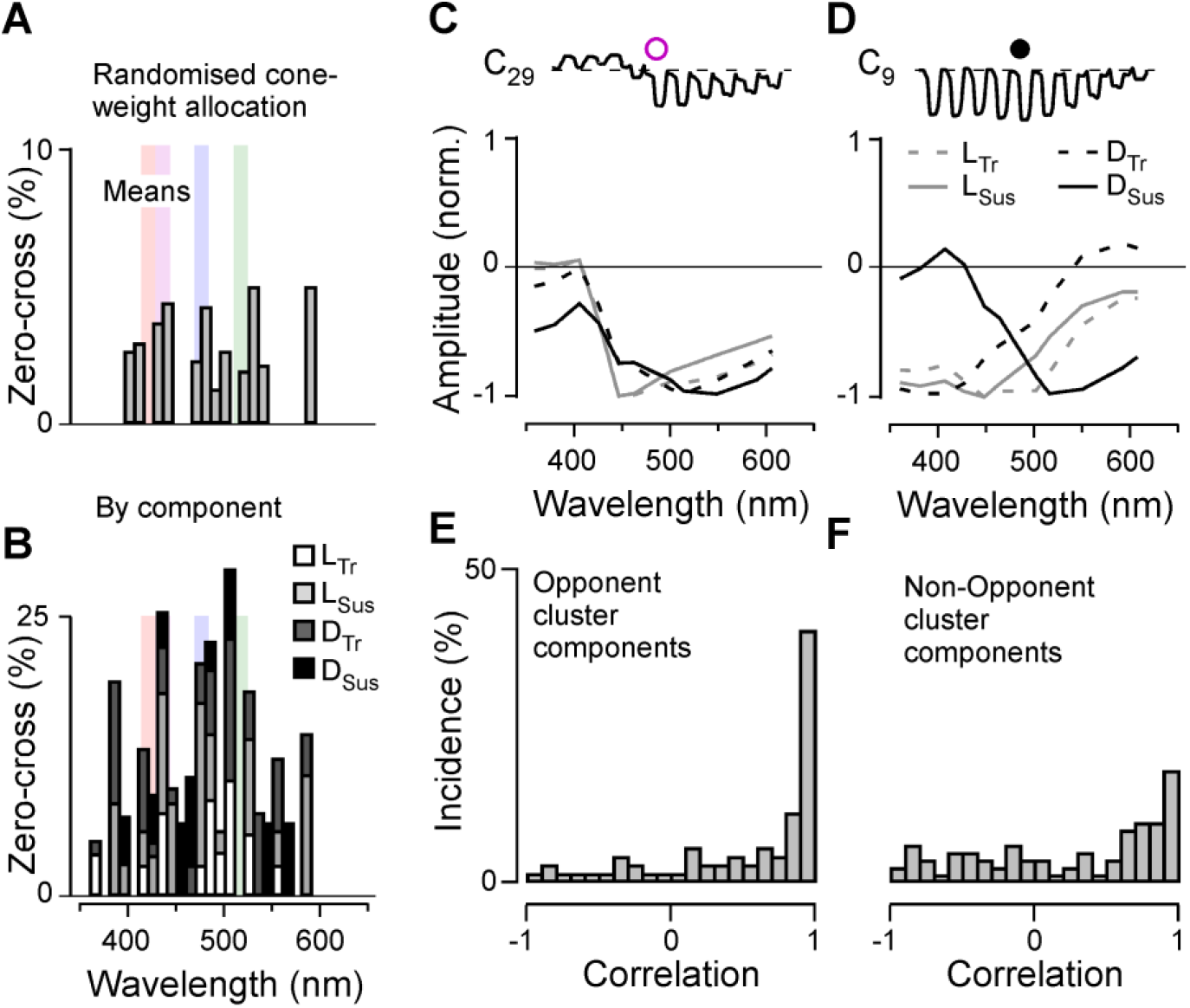
Spectral tunings and temporal components. **A-B**, As Figure 6H,I, but following random permutation of time-components across cones. **C, D**, Spectral tuning functions for two example clusters (C_29_ and C_9_, respectively), computed individually by temporal components as indicated. Note that for C_29_ (C), the four tuning functions were similar to each other, while for C_9_, the tuning of the dark-sustained component deviated strongly from that of the remaining three components. Corresponding time-component resolved tuning functions are detailed for each cluster in Appendix 1. **E**,**F**, Distribution of correlations between each cluster’s “time-component spectral tuning functions” as illustrated in (C,D), for spectrally opponent clusters (E), and for non-opponent clusters (F).

### UV-cone, but not red-/green-/blue-cone weights follow traditional IPL On-Off lamination

Finally, we asked where the inferred new form of UV:R/G/B opponency might be set-up in the inner retina (Figure 7). To this end, we combined the cone-weight data (Figure 4) with information about each BC-terminal’s stratification depth within the inner plexiform layer (IPL) (Figure 3C). In general, the IPL of all vertebrates studied to date is dominated by “Off-circuits” in the upper strata, adjacent to the somata of BCs and most amacrine cells, and by “On-circuits” in the lower strata, adjacent to the somata of retinal ganglion cells^27^. Accordingly, light-components L_tr_ and L_sus_ are expected to mostly exhibit sign-conserving weights in the upper strata, and mostly sign-inverting weights in the lower strata (Figure 7A). Dark components D_tr_ and D_sus_ are expected to exhibit the reverse distribution (Figure 7B).

**Figure 7.**
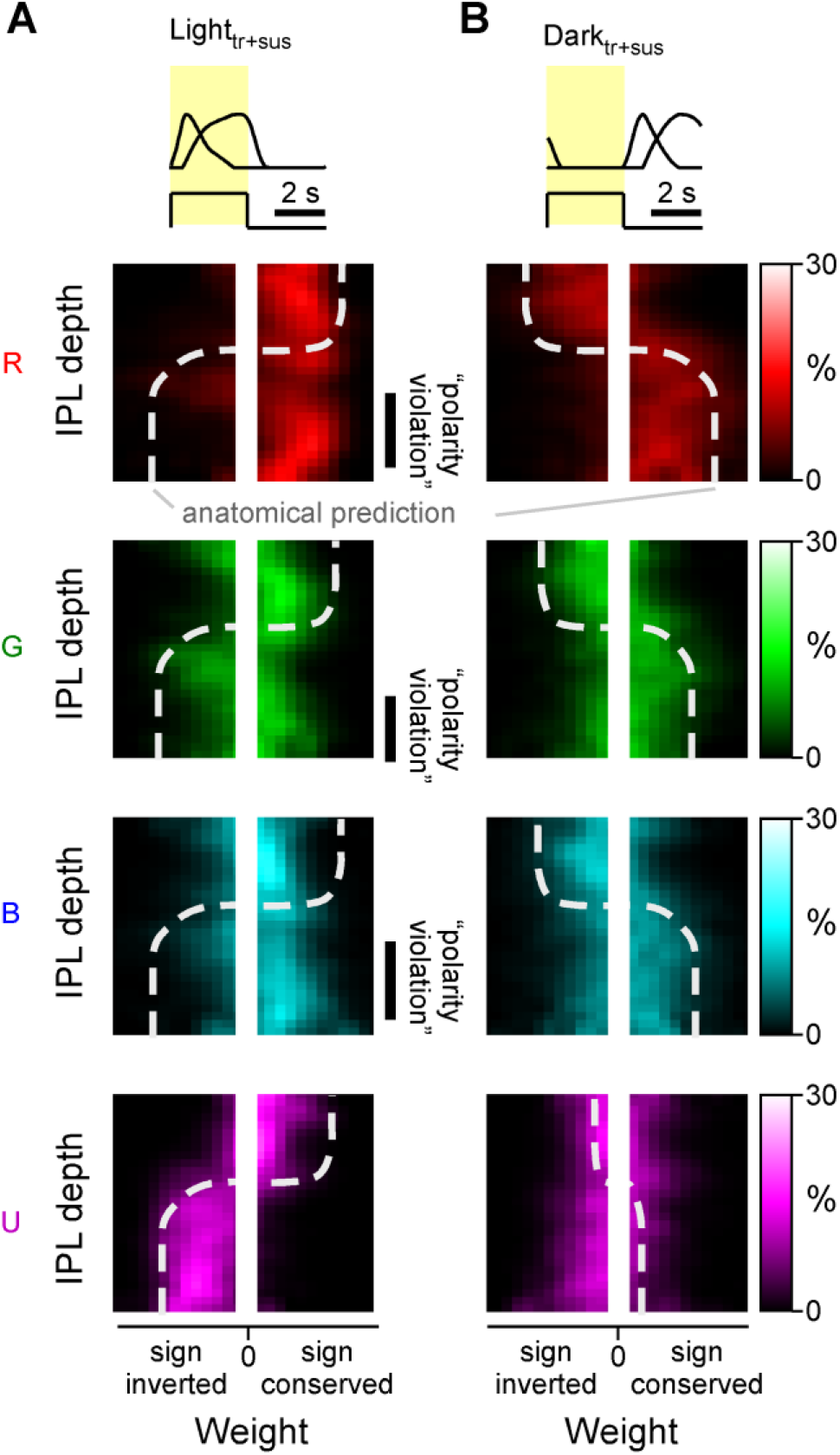
Cone-weight distribution across the inner plexiform layer. **A**,**B**, Two-dimensional histograms of weights (x-axes) associated with each cone resolved by IPL position (y-axes). Brighter colours denote increased abundance. For simplicity, the weights associated with the light (L_tr_, L_sus_) and dark-components (D_tr_, D_sus_), are combined in panels A and B, respectively. Moreover, near-zero weights are not shown (central white bar in all panels). The thick white dotted lines indicate approximate expected distribution of weights based on traditional “On-Off” lamination of the inner retina. By each panel’s side, instances where this expectation is violated are highlighted as “polarity violation”.

This textbook expectation, here graphically indicated by dashed lines, was indeed approximately met when considering dark-components (Figure 7B - note that UV-dark component weights were generally small and not further considered) and for light-components of UV-cones (Figure 7A, bottom panel). Similarly, this classical IPL organisation was also met by red-, green- and blue-cone weights for the upper two-thirds of the IPL, which included the traditional Off-layer, and the upper part of the traditional On-layer (Figure 7A, top three panels). However, specifically for red-, green- and blue-cones, the lower third of the traditional On-layer was dominated by weights of the “wrong” polarity (Figure 7A, top three panels). In agreement, most UV:R/G/B opponent clusters stratified in this lower third of the IPL (Figures 3C,4). Together, this suggests that several of these UV:R/G/B clusters are derived from sign-reversed red-/green-/blue-cone inputs onto “native” UV-On BCs, for example by way of amacrine cells.

## DISCUSSION

We have shown that the substantial spectral and temporal diversity of larval zebrafish BCs (Figures 1,2, cf. Refs^18,29^) can be well-captured by a linear combination of inputs from the four spectral cone-types (Figure 3,4). This in turn allowed us to explore the major functional connectivity rules that govern spectral and temporal widefield signal integration by BCs: We find that red-cones overall provide the dominant input to BCs, often complemented by weaker but same-sign inputs from green- and blue-cones (Figure 5A,C,D). Likely as one consequence, BC pathways do not generally set-up new axes of spectral opponency in the mid-to long-wavelength range. Rather, they mostly either conserve and diversify the two major opponent axes already present in the cones (Figure 6D,E) or establish non-opponent circuits (Figure 6A-C). In contrast, inner retinal UV-cone pathways appear to be organised essentially independently to those of red-, green- and blue-cones (Figure 5E). This leads to the consolidation of a third axis of spectral opponency, contrasting long- and mid-wavelength signals against UV (Figure 6F). This third axis appears to mainly stem from a systematic polarity reversal of inputs from red-, green- and blue-cones onto ‘natively-UV-On’ BCs in the lower IPL (Figure 7A).

### Building spectrally opponent BCs

Because spectral opponency is a prominent feature in larval zebrafish cones^26^, BCs may inherit this property rather than set-up new opponent spectral axes by way of ACs. Indeed, the opponency observed in BC cluster C_15_ could be explained based on weighted but all-sign-conserving inputs from all four cones (Figure 4). However, the full picture may be more complex. For example, like C_15_, cluster C_14_ was also opponent, albeit with a stronger long-wavelength response, and in this case the model used weakly sign-inverted red-cone weights alongside sign-conserved green- and blue-cone weights. In fact, most UV:R/G/B opponent clusters (e.g. C_25-29_) required opposition of long versus short-wavelength cone inputs in the inner retina. This hints that inner retinal circuits may generally use a “mix-and-match” strategy to achieve diverse spectral responses by any available route, rather than strictly adhering to any one strategy. This notion is also tentatively supported by the presence of spectrally diverse amacrine cell circuits in adult zebrafish^22^. More generally, it perhaps remains puzzling how the complex interplay of cone pooling in the outer retina with AC inputs in the inner retina, across 29 highly diverse functional-BC-types which presumably express diverse receptors and ion channels^27^, can ultimately be summarised in an functional wiring logic that for the most part simply sums all four cones, or ‘at best’ opposes a red-/green-/blue-system against UV. Resolving this conceptual conflict will likely require targeted circuit manipulations, for example by comparing BC spectral tunings in the presence and absence of amacrine cell inputs, or after targeted cone-type ablations.

Beyond ‘classical’ opponency, several clusters – both opponent and non-opponent – in addition encoded a notable mixture of spectral and temporal information. Interestingly, several of these clusters appeared to be concentrated around the centre of the IPL (e.g. C_20-25_, Figure 2B,C) – a region which also in mammals has been associated with both transient and sustained processing^34,43–45^. In zebrafish, a mixed time-colour code was previously described for the downstream retinal ganglion cells^21^, which now raises the question to what extent ganglion cells may inherit this property from BCs. Moreover, if and how such information can be differentially read out by downstream circuits and used to inform behaviour remains unknown.

### Three axes of spectral opponency

In principle, the four spectral cone types of larval zebrafish could be functionally wired to for tetrachromatic vision. This would require that all four cone types contribute independently to colour vision. Theory predicts that efficient coding of colour should be based on four channels, an achromatic channel with no zero-crossings on the spectral axis, and three chromatic opponent channels with one, two and three zero-crossings respectively^5,46^. However, such a coding strategy is not essential as demonstrated by the trichromatic visual system of many old-world monkeys which is based on two axes of opponency (“blue-yellow” and “red-green”), each with a single zero crossing. In the present study, we find that among zebrafish BCs, three zero-crossings predominate (Figure 6H,I). Here, the single BC cluster with two zero-crossings (C_7_) did not set-up any notable additional spectral crossings either, but instead crossed once in the ‘blue-cone position’, and once again near the ‘UV-red opponent position’ (Figure 6E). Nevertheless, our findings support the notion that at least at the level of BCs, and under the stimulus conditions used in this study, the zebrafish visual system is capable of supporting tetrachromatic colour vision, as observed behaviourally in goldfish^47^. If and how the larval zebrafish BCs’ axes are preserved, diversified, or even lost in downstream circuits will be important to explore in the future. In this regard, both retinal ganglion cells^20,21^ and brain circuits^20,48^ do carry diverse spectral signals, however beyond a global overview^28^ the nature and distribution of their spectral zero-crossings remain largely unexplored.

### Links with mammalian SWS1:LWS opponency

Of the three spectral axes that dominate the zebrafish inner retina (Figure 6, 8A), those functionally linked with green- (RH2) and blue-cone (SWS2) circuits are unlikely to have a direct counterpart in mammals where these cones-type are lost^1,8^. However, the third axis, formed by functional opposition of UV-cone circuits against red-/green-/blue-cone circuits, may relate to one or multiple of the well-studied mammalian SWS1:LWS opponent circuits^49,50^ (Figure 8B,C).

**Figure 8.**
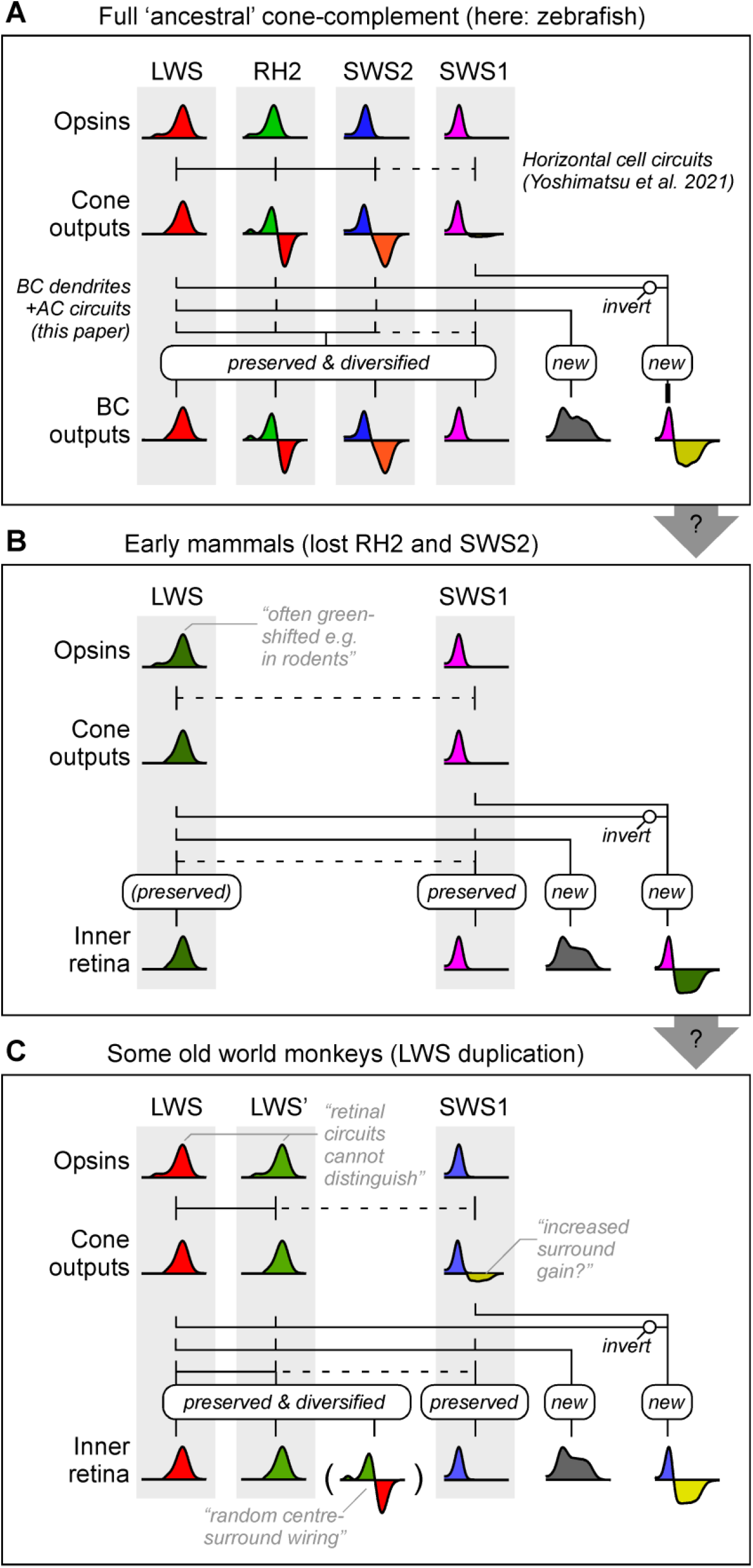
Possible links across vertebrate retinal colour circuits. **A-C**, Conceptual summary schematics of retinal circuits for colour vision in zebrafish (A), dichromatic mammals such as many rodents (B) and some trichromatic old-world monkeys such as humans (C). The coloured ‘graphs’ indicate approximate spectral tuning functions of retinal neurons in a given layer, as indicated.

Despite substantial spectral variation amongst both SWS1 and LWS cone-types across species, mammals usually oppose the signals from SWS1-cones with those of LWS-cones at a retinal circuit level^4,6,51–55^. For example, in the primate outer retina, SWS1-cones exhibit horizontal-cell mediated spectral opponency to LWS signals^56^. Likewise, in the inner retina signals from a highly conserved SWS1-exclusive On-BCs are combined with those of LWS-biased Off-circuits in most if not all mammals that have been studies at this level^35,50,57,58^. Further such circuit motifs can involve diverse but specific types of amacrine and/or retinal ganglion cells^4,53,59^.

Several of these mammalian motifs may have a direct counterpart in zebrafish. For example, like primate SWS1-cones, also zebrafish SWS1-cones exhibit weak but significant long-wavelength opponency that is mediated by horizontal cells^26^. Beyond this possible outer retinal connection, the inferred UV:R/G/B organisation in zebrafish BCs (Figures 5E, 6F, 7) is reminiscent of mammalian circuits associated with SWS1-BCs.

First, as in most mammals^51^, SWS1_On_:LWS_Off_ signals numerically dominate in zebrafish compared to SWS1_Off_:LWS_On_ signals. Second, zebrafish SWS1:LWS opponent signals are predominately found in the lower-most (GCL-adjacent) fraction of the IPL (Figures 3, 7), the same place where mammalian SWS1-On BCs stratify^35^. Third, many zebrafish SWS1_On_:LWS_Off_ signals occurred ventro-temporally (Figure 3D), the retinal region which in mice exhibits the highest density of type-9 BCs^60^, their only SWS1-exclusive BC type^35,57^. While zebrafish are not known to possess an SWS1-exclusive BC^11^, they do possess several anatomical BC types that contact SWS1-cones alongside either one or both of SWS2- (blue) and RH2-cones (green)^8,11^. Such BCs may conceivably become SWS1-exclusive types upon the loss of RH2 and SWS2 cones in early mammalian ancestors.

However, not everything supports a direct correspondence between mammalian and zebrafish SWS1:LWS circuits. For example, in contrast to BCs, among the dendrites of the zebrafish retinal ganglion cells, most UV-opponent signals occur above the IPL midline, near the anatomical border between the traditional On- and Off-layers^21^. Nevertheless, this is approximately in line with the IPL position where several of the well-studied primate SWS1:LWS ganglion cells receive LWS-biased Off-inputs^61^, hinting that similar ganglion cell motifs might also exist in zebrafish. Certainly, zebrafish do possess a number of anatomical retinal ganglion cell types^21,62^ that display similar stratification patterns compared to those that carry SWS1:LWS opponent signals in diverse mammals^50,53^.

A summary of the above argument, showcasing possible links between retinal circuits for colour vision in cone-tetrachromatic species such as zebrafish, to those of most non-primate mammals and of old-world monkeys including humans, is suggested in Figure 8. In the future it will be important to explore if and how mammalian SWS1:LWS circuits can be more directly linked with those found in zebrafish, for example by leveraging molecular markers across potentially homologous types of neurons^36,63,64^.

## METHODS

### RESOURCE AVAILABILITY

#### Lead Contact

Further information and requests for resources and reagents should be directed to and will be fulfilled by the Lead Contact, Tom Baden.

#### Data and Code Availability

Pre-processed functional 2-photon imaging data and associated summary statistics will be made freely available on DataDryad and via the relevant links on http://www.badenlab.org/resources and http://www.retinal-functomics.net.

### EXPERIMENTAL MODEL AND SUBJECT DETAILS

#### Animals

All procedures were performed in accordance with the UK Animals (Scientific Procedures) act 1986 and approved by the animal welfare committee of the University of Sussex. Animals were housed under a standard 14:10 day/night rhythm and fed three times a day. Animals were grown in 0.1 mM 1-phenyl-2-thiourea (Sigma, P7629) from 1 *dpf* to prevent melanogenesis. For all experiments, we used 6-7 days post fertilization (*dpf*) zebrafish (Danio rerio) larvae.

Tg(1.8ctbp2:SyGCaMP7bf) line was generated by injecting pBH-1.8ctbp2-SyjGCaMP7b-pA plasmid into single-cell stage eggs. Injected fish were out-crossed with wild-type fish to screen for founders. Positive progenies were raised to establish transgenic lines. The plasmid was made using the Gateway system (ThermoFisher, 12538120) with combinations of entry and destination plasmids as follows: pBH^65^ and p5E-1.8ctbp, pME-SyjGCaMP7b, p3E-pA^66^. Plasmid p5E-1.8ctbp was generated by inserting a polymerase chain reaction (PCR)-amplified -1.8ctbp fragment^30^ into p5E plasmid and respectively. Plasmid pME-SyjGCaMP7b was generated by replacing GCaMP6f fragment with PCR-amplified jGCaMP7b^67^ in pME-SyGCaMP6f^69^ plasmid.

For 2-photon *in-vivo* imaging, zebrafish larvae were immobilised in 2% low melting point agarose (Fisher Scientific, BP1360-100), placed on a glass coverslip and submerged in fish water. Eye movements were prevented by injection of α-bungarotoxin (1 nL of 2 mg/ml; Tocris, Cat: 2133) into the ocular muscles behind the eye.

### METHOD DETAILS

#### Light Stimulation

With fish mounted on their side with one eye facing upwards towards the objective, light stimulation was delivered as full-field flashes from a spectrally broad liquid waveguide with a low numerical aperture (NA 0.59, 77555 Newport), positioned next to the objective at ∼45°, as described previously^26^. To image different regions in the eye, the fish was rotated each time to best illuminate the relevant patch of photoreceptors given this stimulator-geometry. The other end of the waveguide was positioned behind a collimator-focussing lens complex (Thorlabs, ACL25416U-A, LD4103) which collected the light from a diffraction grating that was illuminated by 13 spectrally distinct light-emitting diodes (LEDs, details below).

An Arduino Due (Arduino) and LED driver (Adafruit TCL5947) were used to control and drive the LEDs, respectively. Each LED could be individually controlled, with brightness defined via 12-bit depth pulse-width-modulation (PWM). To time-separate scanning and stimulating epochs, a global “blanking” signal was used to switch off all LEDs during 2P scanning but enable them during the retrace, at line-rate of 1 kHz (see also Refs^68,70^). The stimulator code is available at https://github.com/BadenLab/HyperspectralStimulator.

LEDs used were: Multicomp Pro: MCL053RHC, Newark: C503B-RAN-CZ0C0AA1, Roithner: B5-435-30S, Broadcom: HLMP-EL1G-130DD, Roithner: LED-545-01, TT Electronics: OVLGC0C6B9, Roithner: LED-490-06, Newark: SSL-LX5093USBC, Roithner: LED450-03, VL430-5-1, LED405-03V, VL380-5-15, XSL-360-5E. Effective LED peak spectra as measured at the sample plane were, respectively (in nm): 655, 635, 622, 592, 550, 516, 501, 464, 448, 427, 407, 381, 360 nm. Their maximal power outputs were, respectively (in µW): 1.31, 1.06, 0.96, 0.62, 1.26, 3.43, 1.47, 0.44, 3.67, 0.91, 0.24, 0.23, 0.20. From here, the first ten LEDs (655 – 427 nm) were adjusted to 0.44 µW, while the three UV-range LEDs were set to a reduced power of 0.2 µW. This relative power reduction in the UV-range was used as a compromise between presenting similar power stimulation across all LEDs, while at the same time ameliorating response-saturation in the UV-range as a result of the UV-cones’ disproportionately high light sensitivity^21,69^. The same strategy was used previously to record from cones^26^.

#### 2-photon calcium imaging

All 2-photon (2P) imaging was performed on a MOM-type 2P microscope (designed by W. Denk, MPI, Martinsried; purchased through Sutter Instruments/Science Products) equipped with a mode-locked Ti:Sapphire laser (Chameleon Vision-S, Coherent) tuned to 927 nm for SyGCaMP6f imaging. We used one fluorescence detection channel (F48×573, AHF/Chroma), and a water immersion objective (W Plan-Apochromat 20x/1,0 DIC M27, Zeiss). For image acquisition, we used custom-written software (ScanM, by M. Mueller, MPI, Martinsried and T. Euler, CIN, Tuebingen) running under IGOR pro 6.3 for Windows (Wavemetrics).

All data was collected using a quasi-simultaneous triplane approach by leveraging an electrically tunable lens (ETL, EL-16-40-TC-20D, Optotune) positioned prior to the scan-mirrors. Rapid axial-jumps of ∼15 µm between scan planes (ETL settling time of <2 ms^31^) were enabled by using a non-telecentric (nTC) optical configuration (nTC_1_, 1.2 mm – see Ref^31^). This nTC optical setup is described in detail elsewhere^31^. All recordings were taken at 128 × 64 pixels/plane at 3 planes (5.2 Hz effective “volume” rate at 1 ms per scan line).

#### Pre-processing of 2-photon data, IPL detection and ROI placement

Raw fluorescence stacks were exported into a Python 3 (Anaconda) environment. The data were de-interleaved and separated into the three recording planes. Next, the data were linearly detrended, linearly interpolated to 42 Hz, and aligned in time. The anatomical borders of the inner plexiform layers were automatically detected by first median-smoothing the time standard deviation images with a Gaussian kernel size of 3 pixels. From here, every pixel above the 35% per-image amplitude threshold was registered as IPL. This automated procedure was made possible by the fact that GCaMP6f expression was restricted to the presynaptic terminals of BCs, which also defined the anatomical borders of the IPL.

To place regions of interest (ROI), a quality index (QI) as described previously^33^ was calculated for each pixel. In short, the QI measures the ratio of variance shared between stimulus repetitions and within a single stimulus repetition. The larger the QI, the more variance in the trace is due to the presented stimulus:

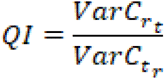

where *C* is the *T* by *R* response matrix (time samples by stimulus repetitions) and × and Var[]x denote the mean and variance across the indicated dimension, respectively. QI ranges from 0 (perfectly random) to 1 (all stimulus repetition responses are identical). This yielded “QI-images” that indicated where in a scan BC-responses were located. From here, ROIs were automatically placed using custom Python scikit-image scripts^71^. In brief, QI-images were adaptively thresholded using kernel size 5. The resulting binary images were distance-transformed and shrunk. The contours of the remaining groups of pixels were recorded and filled, and the highlighted pixels were used as ROI coordinates. From here, the IPL position of each ROI was defined as the relative position of the centre-of-mass of the filled ROI contour to the nearest inner and outer borders of the IPL.

ROI traces were converted to z-scores. For this, a 5 s portion of the trace preceding stimulus presentation was drawn and defined as baseline. The standard deviation of this baseline fluorescence signal was calculated and used to z-score the remainder of the trace. Finally, QIs as described above for each pixel were also calculated for each ROI. ROIs with QI<0.4 were excluded from further analysis. n = 6,125 ROIs passed this quality criterion (72 triplane scans from 7 fish).

#### Clustering of BCs

To identify structure amongst the BC-dataset, trial-averaged ROI traces were PCA-transformed and clustered as described previously (e.g. Refs^18,33^). In brief, we used the first 48 principal components, which accounted for 82% of total variance. Of these, components that near-exclusively carried high-frequency content which is likely linked to noise were discarded. The transformed time-traces were clustered using the scikit-learn (Python 3, Anaconda) implementation of the Gaussian Mixture Models algorithm. The number of clusters was determined using the Bayesian information criterion (BIC). Clusters were judged as stable over repeated clustering runs starting from different random seeds.

#### Reconstruction of BC responses from cones

To reconstruct each BC-mean response into constituent spectral and temporal components, we combined the average spectral tuning curve of each of the four cone-types (from Ref^26^) with four temporal components associated with a given light response (i.e. 3 s On, 3 s Off). The four temporal components used, obtained by non-negative matrix factorisation across all light responses and cluster means, resembled light-transient, light-sustained, dark-transient, and dark-sustained temporal profiles (Figure 3B). Next, each full BC-cluster mean trace was decomposed into a corresponding 4 by 10 array (four temporal components × 10 LEDs; note that we restricted the reconstruction to the central 10 LEDs that generally elicited the greatest variance across BCs). This yielded four spectral tuning curves per BC (i.e. light-transient × 10 LEDs, light-sustained × 10 LEDs and so on), which were then linearly interpolated to the range of 360 - 610 nm to conform with the cone data format. The BC tuning curves were then modelled as linear combinations of the cone tuning curves with a lasso regulariser, which yielded four cone weights X four response bases per BC-trace.

To assess reconstruction quality (Figure S1), reconstructed data was subtracted from the original ROI-means to yield residuals. From here, we compared original data, reconstructions, and residuals by two metrics: variance explained across all clusters, and temporal power explained. To determine the fraction of variance explained by the reconstructions, we first computed the total variance across all clusters for each time-point. The result of this process, plotted beneath each corresponding heatmap (Figure S1A), showed similar time-variance profiles across cluster means and their reconstructions (panels 1 and 2), but very little remaining signal for the residuals (panel 3). From here, we computed the area under the curve for each variance-trace and normalised each to the result from the original cluster means. By this metric, cluster reconstructions captured 94.0% of the original variance, while residuals carried 5.1%.

To determine the extent to which temporal structure was captured, we used a similar approach to the one above, however in this case based on a magnitude-squared Fourier Transform of each time-trace (Figure S1B), limiting the result between 0.08 and 2 Hz which captured the bulk of physiologically meaningful temporal components given the optical imaging approach used (i.e. lower-frequency components would mainly arise from imperfect detrending, while higher-frequency components would exceed the Nyquist recording limit, and further be limited by the kinetics of GCaMP7b. From here, we computed the average of all clusters’ Fourier transforms (plotted beneath each panel) and again computed the faction of this signal captured by the reconstruction (103.8%) and residuals (3.8%). Notably, while this metric was mainly informative about low frequency components which dominated all signals, also higher frequency components were generally well captured, as visible in the individual heatmaps.

### QUANTIFICATION AND STATISTICAL ANALYSIS

#### Statistics

No statistical methods were used to predetermine sample size. Owing to the exploratory nature of our study, we did not use randomization or blinding. To compare weight amplitude distributions (Figure 5A,B) we used the paired Wilcoxon Rank Sum Test, taking paired components as the input (i.e. comparing red-light-transient versus green-light-transient, and so on). To assess weight correlations between cones (Figure 5C-E, Figure S1), we in each case list the Pearson correlation coefficient ρ and 95% confidence intervals (CI) based on the mean weights per cluster. Individual temporal weights were not considered in this analysis. All statistical analysis was performed in Python 3 (Anaconda) and/or Igor Pro 6 (Wavemetrics).

## Acknowledgements

We thank Daniel Osorio and Thomas Euler for critical feedback. The authors would also like to acknowledge support from the FENS-Kavli Network of Excellence and the EMBO YIP.

