## Supplementary figures and images for "Spectral inference reveals principal cone-integration rules of the zebrafish inner retina"

### Appendix 1

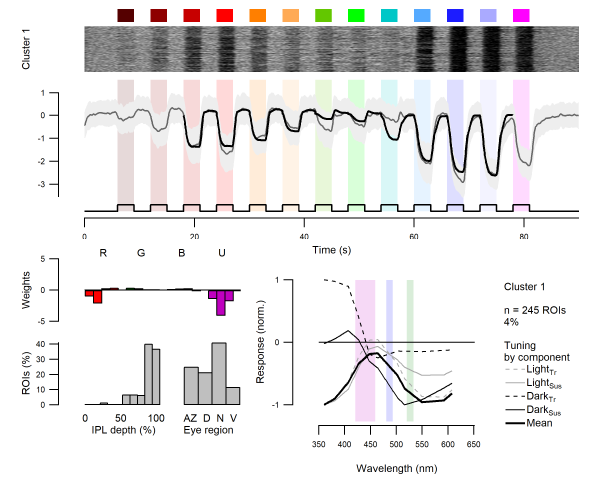
